# Feature overlap modulates rapid semantic but not lexical integration of novel associations by means of fast mapping

**DOI:** 10.1101/594218

**Authors:** Ann-Kathrin Zaiser, Patric Meyer, Regine Bader

## Abstract

There is evidence that rapid integration of novel associations into cortical networks is possible if associations are acquired through a learning procedure called fast mapping (FM). FM requires precise visual discrimination of sometimes highly similar pictures of a previously unknown and a known item, and linking an unfamiliar label to the unknown item. In order to shed light on the mechanisms underlying learning through FM, we manipulated feature overlap between the two items as potential modulating factor. In Experiment 1, we found that labels of the unknown items generally evoked instantaneous lexical competition when encoded through FM, indicating rapid integration into lexical networks. In Experiment 2, we observed semantic priming immediately after FM encoding but only if the items shared many features. This indicates that whereas feature overlap leaves item-level lexical integration unaffected, it might mediate semantic integration of arbitrary picture-label associations, which could explain contradictory findings in the literature.

**Highlights:**

- We examined cortical integration of associations using implicit memory measures.
- Fast mapping enables immediate integration of associations into cortical networks.
- Semantic integration requires the discrimination between items sharing many features.
- Item-level lexical integration is unaffected by feature overlap.

## 1. Introduction

According to traditional theories of declarative memory, consolidation of novel associations is a gradual, time-consuming process. The complementary learning systems theory (McClelland, McNaughton, & O’Reilly, 1995; Norman & O’Reilly, 2003) proposes that storage of new associations into long-term memory underlies two strongly interleaved processes, that is, the initial, fast acquisition of new information by means of hippocampal processing and the gradual incorporation of this information into lexico-semantic networks, represented in cortical structures. Consolidation of novel associations is assumed to be achieved by means of continuous reactivation through hippocampal-neocortical interplay (see Frankland & Bontempi, 2005, for a review).

However, recent findings revealed that rapid, direct integration of novel associations – potentially bypassing the hippocampus – can be successful if a learning procedure called *fast mapping* (FM) is used for knowledge acquisition (e.g., Himmer, Müller, Gais, & Schönauer, 2017; Merhav, Karni, & Gilboa, 2014, 2015; Sharon, Moscovitch, & Gilboa, 2011; see also Coutanche & Koch, 2017, and Coutanche & Thompson-Schill, 2014, for evidence for rapid lexical integration of the labels on an item-level and delayed semantic integration of the associations). In the typical FM paradigm, participants are presented with two pictures of objects, one of which is supposed to be *previously known* (e.g., a flamingo), whereas the other one is supposed to be *previously unknown* (e.g., an exotic, blue-footed bird). Their task is to answer a question referring to a previously unknown label (e.g., *Does the satellote have blue feet?*). In order to do so, participants need to recognize the previously known item, infer that the unknown label refers to the previously unknown item – thereby presumably incidentally creating a picture-label association –, and respond to the question with regard to the unknown item.

Sharon et al. (2011) examined this learning procedure in four amnesic patients suffering from severe lesions to the medial temporal lobe, predominantly to the hippocampus. These patients did not recognize the picture-label associations above chance level if the associations were intentionally learned within a standard explicit encoding (EE) task, in which they were explicitly asked to remember an unknown item together with its label. This might be attributed to their reduction in hippocampal volume as this is a task in which the hippocampus typically would be recruited. Interestingly, when the same patients encoded novel associations within the FM paradigm, their recognition performance was as good as that of healthy controls, implying that hippocampal processing can be bypassed through learning by means of FM^1^.

Despite this evidence that FM might enable successful direct integration of associations, other studies revealed contradictory findings (cf. Greve et al., 2014; Smith et al., 2014; Warren & Duff, 2014; Warren et al., 2016). In order to resolve these contradictions, factors mediating learning success in the FM paradigm yet need to be identified. Sharon et al. (2011) suggested three key determinants to be crucial for successful learning by means of FM: (1) Learning needs to be incidental. (2) The picture-label associations need to be actively discovered by the participants themselves through a process called *disjunctive syllogism*, that is, excluding the previously known item in order to create a link between the label and the unknown item. (3) The new associations need to be learned in the context of previously known information, activating already existing semantic structures into which the new information can be integrated. However, there are studies in which these criteria were entirely fulfilled but still, no learning benefits of FM were observed (e.g., Greve et al., 2014; Smith et al., 2014). Hence, these three determinants might be essential but not necessarily sufficient for successful learning by means of FM and yet undiscovered parameters possibly moderating learning success in the FM paradigm need to be detected.

A promising approach could be to ask which known functional characteristics of candidate brain structures could appropriately support this learning mechanism. Most of the previous literature points to the anterior temporal lobe (ATL), specifically the temporal poles, as a structure critical for FM (e.g., Atir-Sharon, Gilboa, Hazan, Koilis, & Manevitz, 2015; Greve et al., 2014; Merhav et al., 2015; Sharon et al., 2011). This fits nicely with the notion that the ATL serves as an amodal semantic hub, integrating information from modality-specific cortices (see Lambon Ralph, Jefferies, Patterson, & Rogers, 2017; and Patterson, Nestor, & Rogers, 2007, for reviews). It is therefore plausible that these anterior temporal structures may serve as a system supporting rapid semantic integration through FM. Sharon et al. (2011) also reported two patients with additional damage to the ATL who were not able to benefit from FM, which further supports this idea. However, these patients also exhibited extensively reduced volumes of the left perirhinal cortex (PrC). The PrC, a structure located in the anterior part of the medial temporal lobe, was found to be involved in conceptual and perceptual processing of complex, higher-order object representations (e.g., Bussey, Saksida, & Murray, 2005; Cowell, Bussey, & Saksida, 2010; Tyler et al., 2004), specifically in the discrimination of objects sharing many features (e.g., Barense et al., 2005; Bussey, Saksida, & Murray, 2002; Tyler et al., 2013). In addition, the PrC is involved in semantic processing (e.g., Meyer et al., 2013; Meyer, Mecklinger, & Friederici, 2010; Meyer et al., 2005; Wang, Lazzara, Ranganath, Knight, & Yonelinas, 2010; Wang, Ranganath, & Yonelinas, 2014), familiarity-based item recognition (e.g., Bowles et al., 2007; Bowles et al., 2010; Wang et al., 2014; see Brown & Aggleton, 2001, for a review), and also in associative memory but only if item pairs are processed as a single unit (Haskins, Yonelinas, Quamme, & Ranganath, 2008; Quamme, Yonelinas, & Norman, 2007). It is therefore well conceivable that the computational mechanisms of the PrC during the processing of highly complex picture-label associations might be especially qualified to support the encoding and integration of these associations into semantic memory within the FM paradigm. If this is the case, higher demands on perirhinal functions (e.g., object discrimination) during FM encoding might foster integration into neocortical networks through deeper encoding. It is important to note that although higher demands in general can be very resource-consuming and could therefore lead to worse memory, we refer to higher demands selectively on processes involved in FM learning, that is, amongst others, the discrimination of highly complex objects.

Although a key characteristic of the FM paradigm is that the unknown item must be encoded in the context of a previously known item (see Coutanche & Thompson-Schill, 2014, Experiment 2), inter-item similarity has not yet been explicitly manipulated. However, it is noticeable how similar the unknown and the known items were in studies conducted by Sharon and colleagues (2011; see also Sharon, 2010, for further examples). Following the rationale outlined above, we predict that higher feature overlap between the known and the unknown item in the FM task promotes faster and better neocortical integration. As explicit recognition tests do not necessarily measure neocortical integration in healthy young adults but could as well reflect hippocampal reactivation, we tested this prediction using implicit tests. Although conclusions on underlying neurocognitive mechanisms cannot be drawn by means of behavioral measures only, these implicit measures of memory can serve as indicators for (cortical) integration instead of hippocampus-based retrieval as they provide direct access to semantic networks represented in cortical structures and it has been shown that they less likely involve hippocampal processing (Goshen-Gottstein, Moscovitch, & Melo, 2000). In order to implicitly measure integration into lexico-semantic networks, we assessed the effects of the newly learned associations on the processing of already known lexically or semantically related items following a procedure used by Coutanche and Thompson-Schill (2014; see also Coutanche & Koch, 2017). They argued that successful integration of new associations into neocortical structures should result in lexical competition on one hand, and in semantic priming on the other hand. Generally, lexical competition leads to *inhibition* due to interference caused by co-activation of *lexically* neighboring items at retrieval. Therefore, it takes more time until a target word is uniquely identified if it has more lexical neighbors (e.g., slowed response times to *mouse* as it has many lexical neighbors such as *house, horse*, etc.). In contrast, in semantic priming, access to *semantically* related items is *facilitated* (e.g., faster response times to *mouse* if it was preceded by *hamster*). Thus, if new information, such as newly learned labels, lexically competes with or semantically primes old information, it can be assumed that it is integrated into neocortical lexico-semantic networks. Coutanche and Thompson-Schill (2014) found lexical competition, that is, slowed responses to English words which lexically neighbored labels of the learning phase, 10 minutes after encoding trough FM and again after 24 hours. For the EE group, no lexical competition was observed, neither immediately nor on the following day, indicating that successful rapid and persistent lexical integration is possible after encoding through FM but not through EE. In order to measure semantic integration, Coutanche and Thompson-Schill (2014) conducted a semantic priming task, in which newly learned labels of previously unknown animals were expected to prime semantically related but not unrelated targets. Contrary to their expectations, no priming effects were found for neither encoding condition after 10 minutes. After 24 hours, they found a significant priming effect for the FM group only. Unfortunately, potential confounds could have influenced the data pattern. As related targets in the priming phase were always animals and unrelated targets were always artifacts, semantic categories of the targets were not counterbalanced. Since response latencies between related and unrelated targets might have differed already on a baseline level, it cannot be excluded that faster processing of the artifacts could have masked a potential priming effect. In addition, semantic integration was measured using a lexical decision task, which might not have required sufficient semantic processing in order to observe semantic priming effects (see De Houwer, Hermans, Rothermund, & Wentura, 2002).

In order to investigate the role of feature overlap in FM learning, we followed Coutanche and Thompson-Schill’s (2014) experimental design, but with a few important adaptations. In Experiment 1, we examined lexical integration by means of FM separately for a condition in which the previously unknown and the known item share many features (*fast mapping, high overlap*; FMHO) compared to a condition in which they share few features (*fast mapping, low overlap*; FMLO), assuming that rapid integration into lexico-semantic networks can be fostered by a high feature overlap between the previously unknown and the known item at encoding. In Experiment 2, we investigated if immediate semantic integration by means of FM is possible and if it can be strengthened in an FMHO condition but not in an FMLO and an EE condition. We assessed semantic integration both immediately after encoding and again after 24 hours in order to examine stability over time.

## 2. Experiment 1

Analogously to Coutanche and Thompson-Schill (2014), we used a lexical competition task in order to measure lexical integration. For this purpose, labels in the encoding task needed to be artificially created lexical neighbors of already existing English hermit words (i.e., words which are not transformable into other words by changing one letter). If such hermit words that naturally do not have any lexical neighbors (e.g., *tomato*) obtain a new lexical neighbor at encoding (e.g., if the label *torato* is successfully learned), the relative increase of the number of neighbors of the hermits is large. Therefore, competition effects are expected for responses to hermits that obtained a new neighbor at encoding but only if this new neighbor has been successfully integrated. We expected to observe a general lexical competition effect for associations acquired by means of FM. This competition effect was assumed to be larger when the known and the unknown item share many features (FMHO) than when they share few features (FMLO). As stable lexical competition effects for FM and no effects for EE have previously been reported (Coutanche & Thompson-Schill, 2014), we decided to only set focus on the effects of feature overlap within the FM paradigm in Experiment 1. In addition to this implicit measure of integration, we conducted a forced-choice recognition test in order to examine if participants also showed explicit learning above chance level. As it cannot be disentangled if recognition accuracy in healthy young participants is driven by hippocampus-dependent retrieval or by retrieval of associations already incorporated into lexico-semantic networks, we did not make assumptions about differences in recognition accuracy between the overlap conditions. However, assessing recognition accuracy was necessary in order to tell if a potential lack of a lexical competition effect would have been an issue of encoding difficulties (e.g., too short presentation times, too difficult questions, etc.) or if selectively rapid neocortical integration did not work but there still was explicit (perhaps hippocampal) learning.

### 2.1 Methods

#### 2.1.1 Participants

Thirty-six students from Saarland University took part in the experiment (*M*_age_ = 23.4 years, age range: 20-30 years; 31 female). All participants were native German speakers and had normal or corrected-to-normal vision. Participants gave written informed consent prior to the experiment and completed the experiment within approximately 50-60 minutes. They were compensated for their participation with 8€ per hour. The experiment was approved by the local ethics committee of Saarland University in accordance with the declaration of Helsinki.

#### 2.1.2 Materials

All pictures were obtained from the internet and were drawn from an item pool of a previously conducted rating study, in which a different sample of 46 participants (*M*_age_ = 23.1 years, age range: 18-34 years; 30 female) had rated pictures of items of eight categories (mammals, birds, insects, fish, reptiles, fruit, vegetables, plants) for familiarity (5-point Likert scale; 1 = *not at all familiar*, 5 = *very familiar*) and previous knowledge (*known* vs. *unknown*). At stimulus creation for the rating study, each of 180 putatively unknown items was assigned two putatively known items, thereby creating two picture pairs per unknown item (see Figure 1). One of the putatively known items was supposed to be highly similar to the unknown item (for usage in the FMHO condition) and the other one less similar to the unknown item (for usage in the FMLO condition). The putatively known items were also used with two different unknown items, such that a putatively known item (e.g., a flamingo) was once part of a high-overlap item pair together with an unknown, similar-looking item (e.g., a wading bird) and once part of a low-overlap item pair together with another, dissimilar unknown item (e.g., a mouse-like mammal). Putatively known items of such a pair of item pairs were interchanged for the respective other unknown item (e.g., guinea pig as low-overlap known item for the wading bird and as high-overlap known item for the mouse-like mammal). Each participant was presented with either the high-overlap or the low-overlap item pair (counterbalanced between subjects). Feature overlap between the two pictures of an item pair, which was defined as the number of features the two pictures have in common, was rated on a 5-point Likert scale (1 = *not at all similar*, 5 = *very similar*). Examples in the instructions of the rating study made clear that feature overlap refers to features such as the presence and nature of fur, a tail, a fin, legs, the smoothness of a fruit’s skin, color, and so forth.

**Figure 1.**
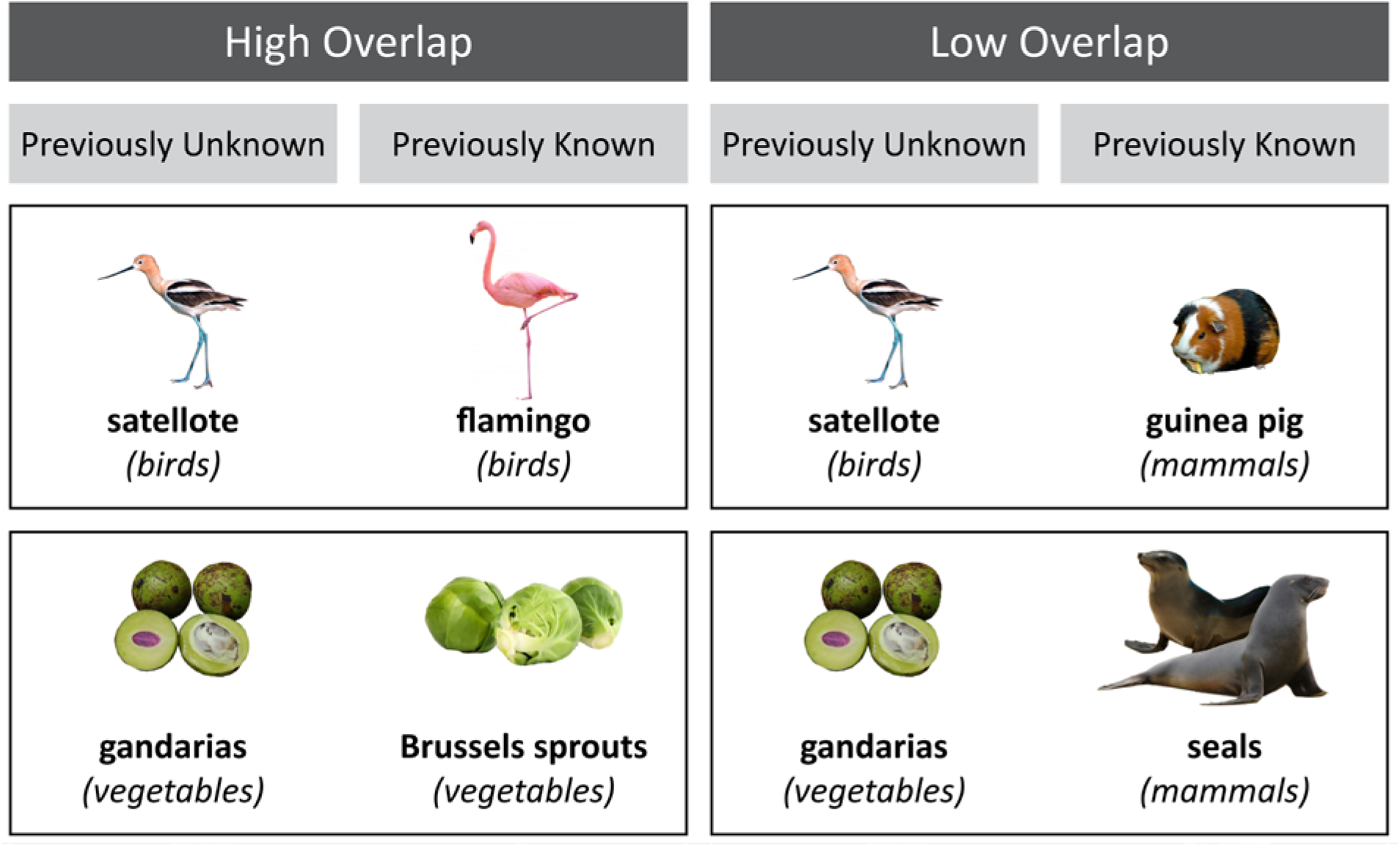
Example item pairs. Each line depicts a pair of item pairs, consisting of one previously unknown item and two previously known items (one for each overlap condition). Low-overlap item pairs could consist of items from the same higher-level category (upper line: both animals, with *bird* and *mammal* as lower-level categories) or from different higher-level categories (lower line: plants and animals, with *vegetables* and *mammals* as lower-level categories).

The encoding phase in Experiment 1 contained 92 pairs of previously known and unknown items, arranged in two lists, which were assigned to one half of the sample each. In each list, 46 item pairs were presented in the FMHO encoding condition and 46 pairs in the FMLO condition (counterbalanced between lists). Lists did not differ with regard to feature overlap ratings between participants, neither for FMHO trials nor for FMLO trials (both *t*s < 1). Between overlap conditions, semantic categories of the items were distributed equally and items did not differ with regard to familiarity ratings or ratings of previous knowledge, neither of the previously known nor the unknown items (all *p*s > .219). Within each overlap condition, 50% of the questions at encoding required a positive response, 50% a negative response, and the question referring to a previously unknown item was identical for both overlap conditions. Only those item pairs were included for which the previously unknown item had been classified as unknown by most participants in the rating study (on average, by 87 %; *SD* = 12 %) and had been rated with the lowest familiarity (*M* = 2.09, *SD* = 0.45), and for which the previously known item had been rated as known by most participants in the rating study (on average, by 91 %, *SD* = 12 %) and with the highest familiarity (*M* = 4.41, *SD* = 0.39). Moreover, only pairs of item pairs with the highest difference between the overlap ratings of the high-overlap item pair (e.g., satellote – flamingo; see Figure 1) and the low-overlap item pair (e.g. satellote – guinea pig) were included (*M*_FMHO_ = 3.57, *SD*_FMHO_ = 0.49; *M*_FMLO_ = 1.41, *SD*_FMLO_ = 0.32; *M*_diff_ = 2.16, *SD*_diff_ = 0.56). In the final item set, familiarity for the previously unknown items was significantly lower than for the previously known items, significantly more participants of the rating study had rated the previously known item as known than the previously unknown items, and overlap of the high feature overlap pairs was higher than overlap of the low feature overlap pairs (all *p*s < .001). In addition, the high overlap item pair with the lowest overlap still had a higher overlap rating than the low overlap item pair with the highest overlap.

Additional 20 item pairs (10 FMHO, 10 FMLO) were added as filler trials, in which the question referred to the previously known item in order to prevent participants from always responding with regard to the unknown item without paying attention to the label. In order to prevent primacy and recency effects, further two filler trials were inserted as buffer trials each at the beginning and at the end of the encoding phase. Filler trials were excluded from all analyses. The size of all pictures in the experiment varied depending on their relative size in reality, but was 300 × 300 pixels at maximum, leading to a maximum visual angle of approximately 8.2°.

In order to measure lexical competition, we created 48 new lexical neighbors to existing concrete German nouns (see Appendix). We will refer to the latter as hermits, albeit eleven of them already had one lexical neighbor (but with a mean normalized lemma frequency of the neighbor of < 0.01 per million words, *SD* < 0.01; Dudenredaktion, 2009; Heister et al., 2011). Word length of the hermits was between 4 and 8 letters (*M* = 6.50, *SD* = 0.98) and normalized lemma frequencies between 0.52 and 133.94 per million words (*M* = 19.58, *SD* = 34.25; Heister et al., 2011). The artificially created new labels should deviate from the hermit words in one phoneme at maximum, either by adding, deleting, or substituting a phoneme, and this deviation should preferably occur late in the word, in order to shift the point of uniqueness backwards and thus provoke maximum lexical competition with the hermits (Davis & Gaskell, 2009; Gaskell & Dumay, 2003). Of the 48 newly created labels, 32 were used as labels in the encoding phase (16 within FMHO trials, 16 within FMLO trials) and their respective hermits were later used as *neighbor hermits* in the lexical competition task (e.g., *satellite* tested as neighbor hermit if the label *satellote* was encoded). The remaining 16 labels were not encoded as their direct lexical neighbors were used as *non-neighbor hermits* in the lexical competition task (e.g., *satellite* as non-neighbor hermit if the label *satellote* was not encoded). The allocation of labels to neighbor hermit FMHO trials, neighbor hermit FMLO trials, or non-neighbor hermit trials was counterbalanced between subjects, which required that each item was assigned three labels, with each appearing together with this item in one third of the participants. Labels of the remaining 60 items that were not used for the lexical competition task were substituted either with a pseudo-word or with an item’s botanical or zoological name (sometimes slightly modified) if these labels might have subjectively triggered expectations about an item’s category or features. For example, items were renamed if parts of the name included information about the category, such that *giraffe gazelle* (which was given its alternative name *gerenuk* in our experiments) would indicate that the item is an animal. Word length of all labels, including the newly created neighbors of the hermits, was between 4 and 10 letters (*M* = 7.21, *SD* = 1.17).

Each test display in the two-alternative forced-choice recognition test consisted of a label used in the encoding phase, its respective associated picture, and one foil picture. Test foils had all been used as previously unknown items in the encoding phase in order to control for item familiarity. Moreover, both pictures of a test screen were from the same higher-level category: They were either both plants or both animals. Thus, it was not sufficient to remember an item’s semantic category but participants were required to retrieve the specific picture-label combination. No two test pictures appeared together twice. In order to prevent participants from developing strategies, additional 12 filler trials were included, in which both pictures had already been presented twice.

#### 2.1.3 Design and Procedure

Stimulus presentation and timing were controlled using the experimental software PsychoPy (Peirce, 2009; http://www.psychopy.org/). Participants were seated in front of a 17-inch screen, at a viewing distance of approximately 60 cm. All stimuli throughout the experiment were presented against a white background.

##### Encoding

In order to ensure that encoding was incidental, participants were told that the experiment aimed to investigate visual perception of animals, fruit, vegetables, and plants. All participants encoded the associations by means of FM, and feature overlap was manipulated within subjects. Participants first completed six practice trials (3 FMHO, 3 FMLO), followed by the 116 experimental trials (including 24 filler and buffer trials), which were presented in random order with the constraint that stimulus presentation began and ended with two filler trials each. At the beginning of each trial, a fixation cross was displayed for 700 ms, horizontally centered and slightly below the center of the screen, at the same height as the question would appear. The question was then displayed for 5500 ms, with the plain text presented separately for the first 2000 ms (Arial 27 point font) and together with the pictures for 3500 ms (see Figure 2). The label within the question was always presented in the horizontal center of the screen and in bold font. Participants were encouraged to read the question thoroughly, focus on what exactly is asked for and, as soon as the pictures appear, to figure out to which item the question refers and how it is thus to be answered. After the pictures and the question had disappeared, the words *yes* and *no* were displayed in orange and blue color on the left and right side of the screen (color and position counterbalanced between subjects), requesting to press the key marked with the respective color on a computer keyboard. As soon as an answer was given, participants received feedback and the next trial started. If no answer was given within 3000 ms, they were encouraged to respond faster and moved on to the next trial.

**Figure 2.**
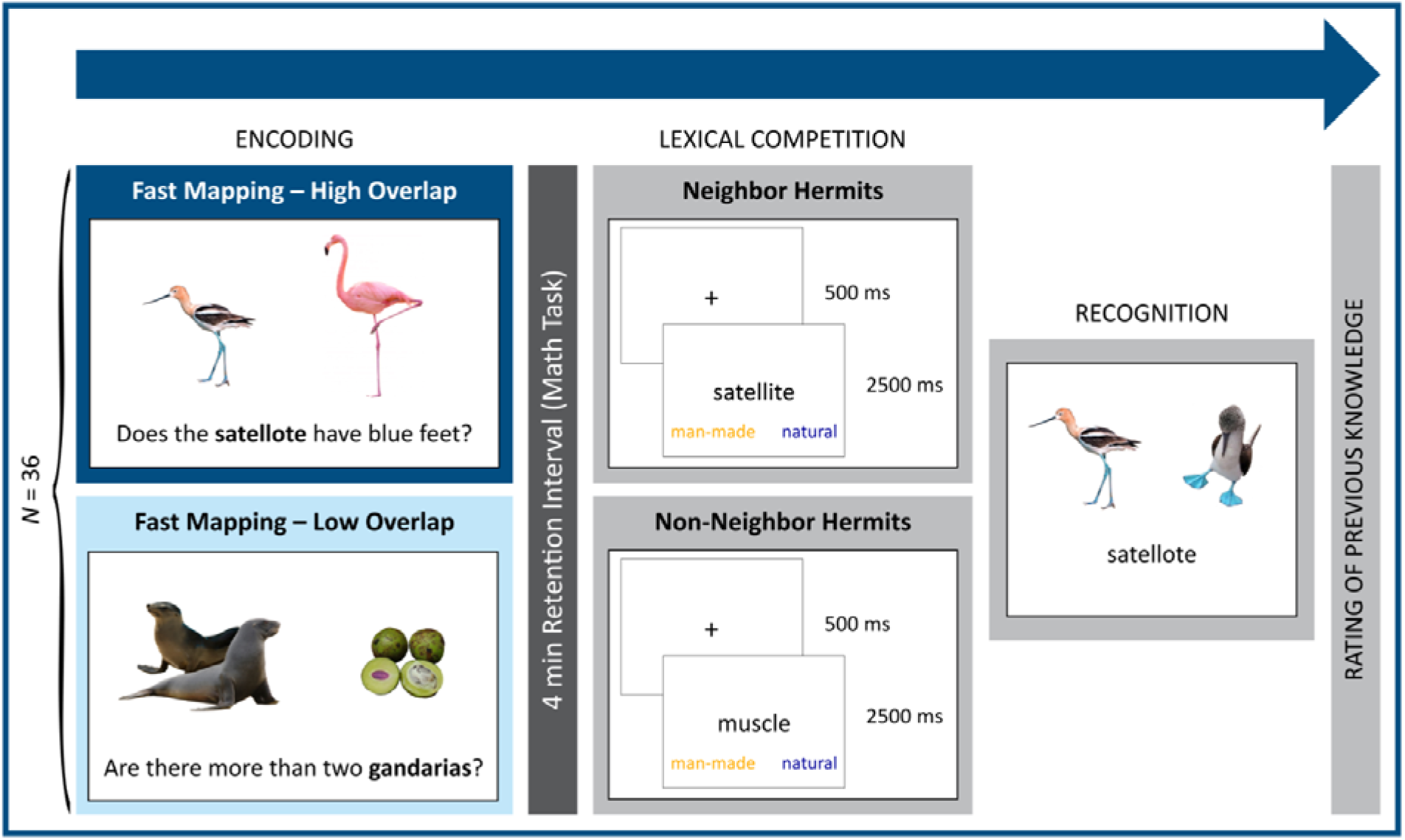
Experimental design and procedure of Experiment 1. **Encoding.** Encoding condition was manipulated within subjects. After the question had been presented for 2000 ms, pictures were inserted and presented together with the question for 3500 ms. Response options (*yes*/*no*) were provided after both pictures and question had disappeared. Of 92 unknown items, 32 were renamed in order to serve as new lexical neighbors for the lexical competition task (e.g., *satellote* as a neighbor for the hermit *satellite*). Feedback was given after a response had been made. **Lexical competition.** In the lexical competition task, responses were given to 32 hermits that had obtained a new neighbor at encoding (neighbor hermits) versus 16 hermits that had not obtained a new neighbor (non-neighbor hermits). Sixteen lexical neighbors of the 32 hermits were encoded in the FMHO (fast mapping, high overlap) condition and 16 in the FMLO (fast mapping, low overlap) condition. **Recognition.** In the two-alternative forced-choice recognition test, targets and foils within one display always belonged to the same higher-level category (i.e., either both items were animals or both items were plants).

##### Lexical competition

After a retention interval of 4 minutes, in which participants solved simple mathematical equations, the lexical competition phase was administered. First, participants were familiarized with the task in a practice phase consisting of four trials using German nouns that did not appear elsewhere in the experiment. In contrast to the experimental trials, feedback was given at the end of each practice trial and participants were encouraged to respond faster if they had not responded within the given time window of 2500 ms. The actual lexical competition task contained 48 trials, which were presented in random order. Each trial began with a fixation cross in the center of the screen for 500 ms, followed by the presentation of the hermit word (see Figure 2). Hermits were displayed in Arial 27 point font. Participants were instructed to decide if a hermit is man-made or natural by keypress. The words *man-made* and *natural* were displayed in blue and orange color on the left and right side on the bottom of the screen (color and position counterbalanced between subjects). The next trial started as soon as a response was given but after 2500 ms of stimulus presentation at maximum. Instructions emphasized speed over accuracy and participants were additionally informed that due to the fast pace of the task, they might make mistakes but nevertheless should focus on responding as fast as possible (as recommended by Wentura & Degner, 2010).

##### Recognition

In the recognition test, participants were presented with two pictures and a label, and were asked to indicate which of the pictures belonged to the label. After the presentation of a fixation cross for 500 ms in the center of the screen, the label was displayed horizontally centered slightly underneath the position of the fixation cross in Arial 27 point font, together with a test target and a test foil picture to the left and to the right (50% of the target and foil pictures on each side) slightly above the position of the fixation cross (see Figure 2). This test display stayed on the screen until a response was made by pressing the respective left or right key on the computer keyboard, but for 3500 ms at maximum. If no key had been pressed within this time, participants were encouraged to respond faster and the next trial started. All 92 picture-label associations were tested, including the 32 associations of which the neighbor hermits were presented in the lexical competition task. Each picture of an unknown item was presented twice, once as target and once as foil. Twenty-four unknown pictures were presented three times in order to create 12 additional filler trials. Repeated presentations of a picture were separated by at least eight trials and no combination of test pictures appeared twice. Both pictures of a test display were encoded within the same encoding condition. Again, this phase was also preceded by a practice phase, in which the items from the encoding practice phase were used as test items. Feedback was given only in the practice phase.

##### Rating of previous knowledge

At last, previous knowledge of all items was assessed with a rating scale. After debriefing participants about the intention of the study and the renaming of the stimuli, they were instructed to indicate on a 5-point Likert scale how well they had known the item before the experiment, no matter under which name (1 = *had not known the item at all before the experiment*; 5 = *had known the item very well before the experiment*). After ratings of >= 4, participants were asked to type in the item’s name at the lowest category level possible (e.g., *hawk* instead of *bird*).

#### 2.1.4 Data analysis

All analyses were conducted using R (R Core Team, 2016). Lexical competition effects were calculated by subtracting response times for correct responses to non-neighbor hermits from response times for correct responses to neighbor hermits. Trials were removed if they contained items for which a participant’s individual rating of prior knowledge was inconsistent with the expected knowledge (rating of <= 3 for previously known items and >= 4 for previously unknown items; mean dropout rate: 5.7 % of correct trials). We further excluded outlier trials with regard to response times individually for each participant according to the outlier criterion recommended by Tukey (1977; 1.5 inter-quartile ranges below the first and above the third quartile) and, in line with Coutanche and Thomson-Schill (2014; see also Bowers, Davis, & Hanley, 2005) and Coutanche and Koch (2017), all trials with response latencies below 300 ms and above 1500 ms as too long response times are unlikely to be influenced by implicit processes. This resulted in a final mean dropout rate of 12.6 % of correct trials. There were no outlier participants (Tukey, 1977) regarding the lexical competition effect. Recognition accuracy represents the proportion of correct responses. If not noted differently, *t* tests to compare lexical competition effects were one-tailed and the significance level was set to α = .05. Effect size *d* for the within-subjects comparison of the lexical competition effect were calculated as difference of the mean lexical competition effects divided by the pooled standard deviation of the difference and corrected for the within-subjects correlation of the effects (see Morris & DeShon, 2002). Effect size *d* for the between-subjects deviation of the lexical competition effect from zero was calculated as the mean lexical competition effect divided by the standard deviation of the effect. Effect size *d* for the between-subjects deviation of recognition accuracy from zero was calculated as the mean recognition accuracy divided by the standard deviation of recognition accuracy.

### 2.2 Results

#### Lexical competition

All participants performed above chance level in the lexical competition task (*p* < .05, binomial test; see Table 1 for accuracies). The accuracy difference between neighbor hermits and non-neighbor hermits was only marginally significant, *t*(35) = −1.99, *p* = .054, and neither reached significance for the FMHO condition, *t*(35) = 1.78, *p* = .084, nor for the FMLO condition *t*(35) = 1.75, *p* = .090, all two-tailed. Although we only observed a lexical competition effect for the FMLO condition, *t*(35) = 2.02, *p* = .025, *d* = 0.34, but not for the FMHO condition, *t*(35) = 1.10, *p* = .141, there was a general lexical competition effect, *t*(35) = 1.94, *p* = .030, *d* = 0.33 (mean competition effect: *M* = 16.36 ms, *SD* = 50.69 ms; see Figure 3), that is, response times to neighbor hermits were significantly slower compared to non-neighbor hermits (see Table 1 for response times). Lexical competition in the FMHO condition (mean competition effect: *M* = 11.24 ms, *SD* = 61.56 ms) was numerically smaller than in the FMLO condition (mean competition effect: *M* = 20.25 ms, *SD* = 60.08 ms), contrary to our hypotheses. However, exploratory post-hoc analyses revealed that the lexical competition effect was not significantly different between FMHO and FMLO trials, *t*(35) = 0.81, *p* = .423, two-tailed.

**Table 1.**
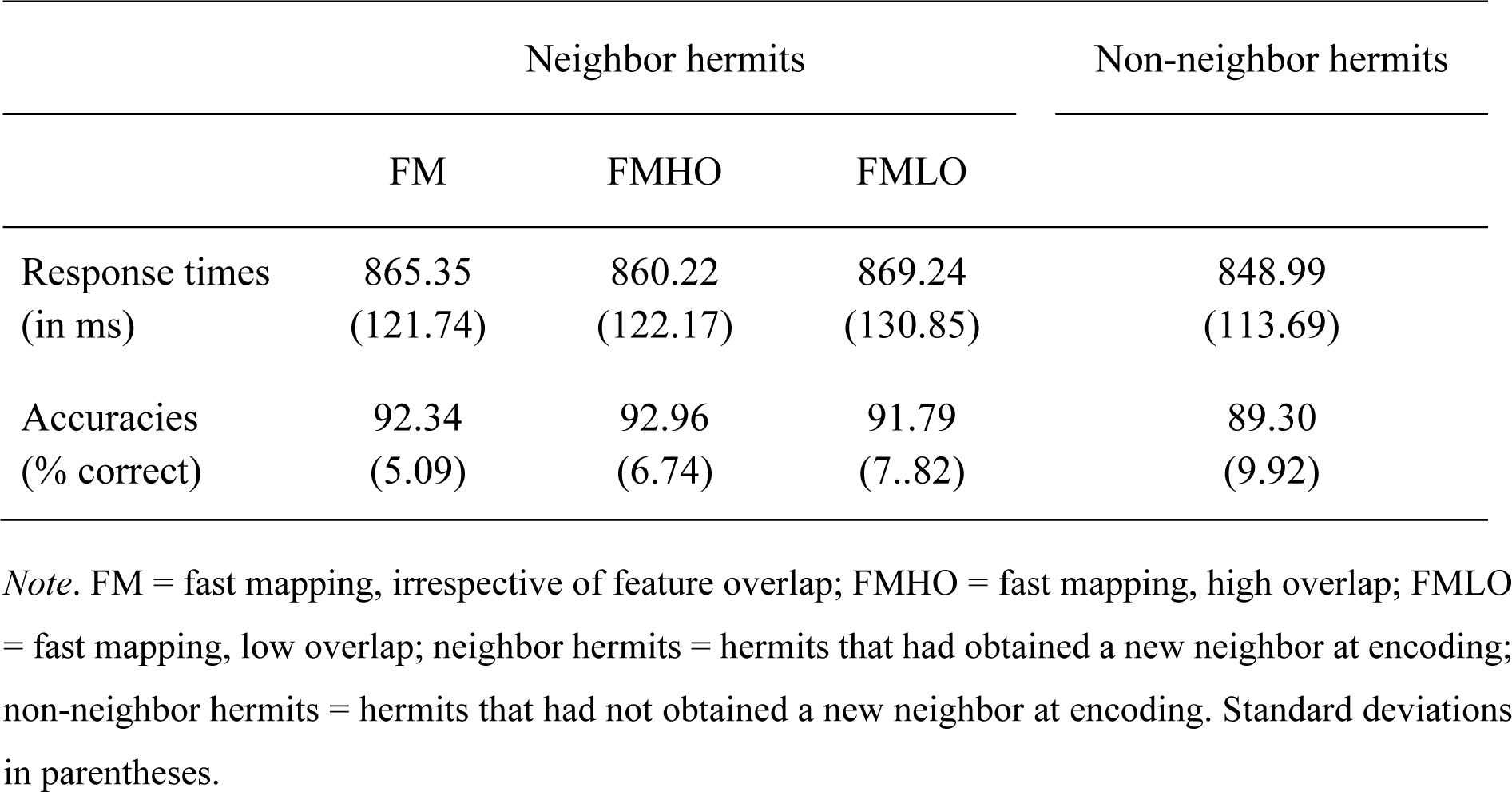
Mean Response Times (in ms) and Accuracies (in % Correct) to Neighbor Hermits and to Non-Neighbor Hermits by Encoding Condition in the Lexical Competition Task of Experiment 1

|  | Neighbor hermits |  |  | Non-neighbor hermits |
| --- | --- | --- | --- | --- |
|  | FM | FMHO | FMLO |  |
| Response times<br>(in ms) | 865.35<br>(121.74) | 860.22<br>(122.17) | 869.24<br>(130.85) | 848.99<br>(113.69) |
| Accuracies<br>(% correct) | 92.34<br>(5.09) | 92.96<br>(6.74) | 91.79<br>(7..82) | 89.30<br>(9.92) |

*Note.* FM = fast mapping, irrespective of feature overlap; FMHO = fast mapping, high overlap; FMLO = fast mapping, low overlap; neighbor hermits = hermits that had obtained a new neighbor at encoding; non-neighbor hermits = hermits that had not obtained a new neighbor at encoding. Standard deviations in parentheses.

**Figure 3.**
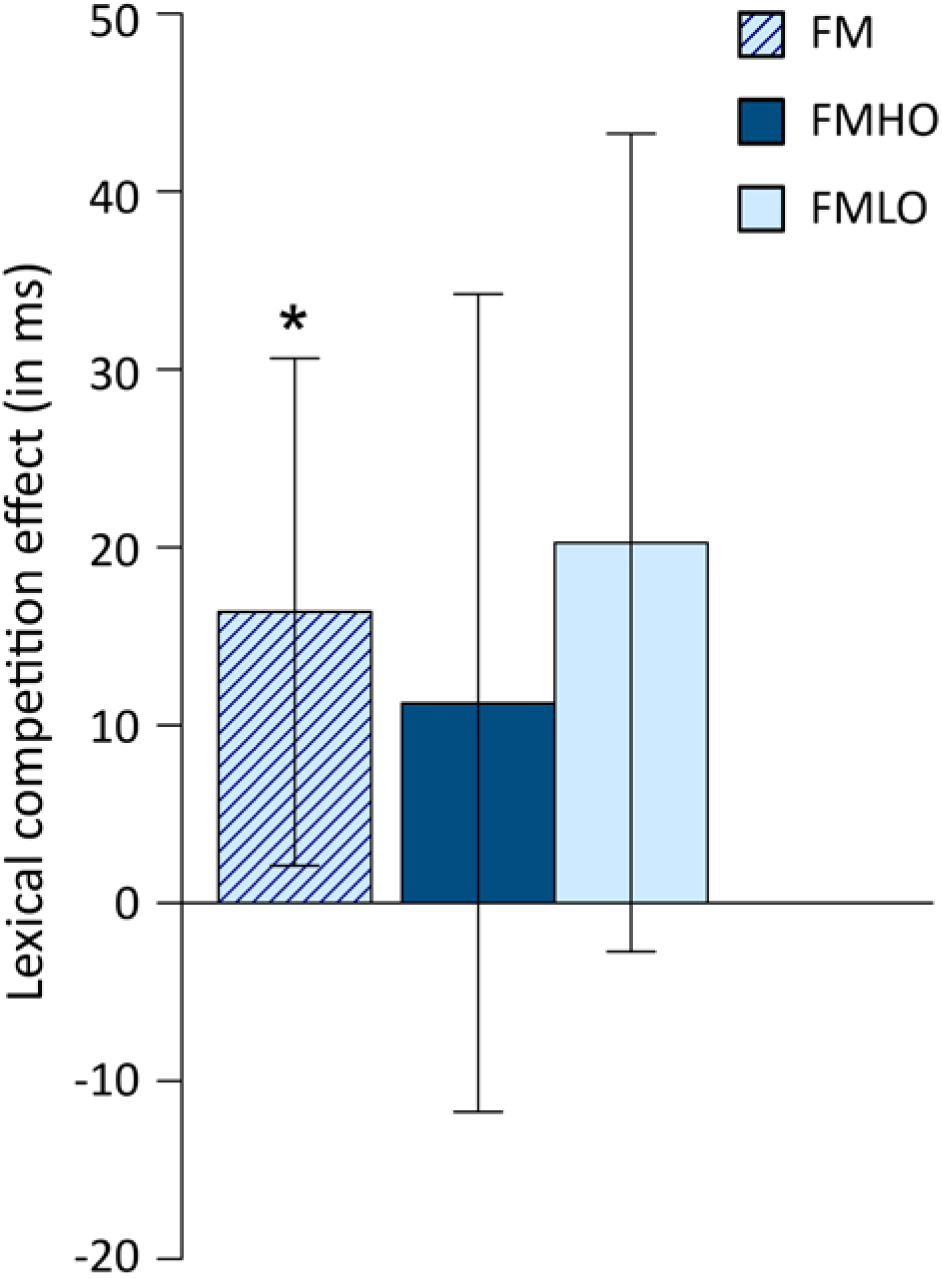
Results of the lexical competition task of Experiment 1. The lexical competition effect was calculated by subtracting response times for responses to words that had not obtained a new neighbor at encoding (non-neighbor hermits) from response times for responses to words that had obtained a new neighbor (neighbor hermits). FM = fast mapping, irrespective of feature overlap; FMHO = fast mapping, high overlap; FMLO = fast mapping, low overlap. Error bars for the FM condition represent the one-tailed confidence interval for the lexical competition effect. Error bars for the FMHO and FMLO conditions represent the two-tailed within-subject confidence interval for the differences between the lexical competition effect in the FMHO condition and in the FMLO condition. \**p* < .05

Although the pattern of the accuracy data (see Table 1) might indicate a tendency towards a speed-accuracy trade-off, differences in accuracies, which could reflect such a trade-off, did not reach significance. In order to further investigate if a lexical competition effect is also apparent in a sample with an accuracy pattern contrary to what would indicate a speed-accuracy trade-off, we examined lexical competition in a subgroup of participants who showed numerically higher accuracies for non-neighbor hermits than for neighbor hermits. In this group (*N* = 16), a lexical competition effect was also found, *t*(15) = −1.85, *p* = .042, *d* = 0.46, one-tailed, indicating that even if a speed-accuracy trade-off could definitely be excluded, there still was rapid lexical integration.

#### Recognition

In order to investigate whether participants also showed above-chance explicit associative memory, we checked accuracy in the recognition test. Participants performed above chance level in the FMHO condition, *t*(35) = 4.25, *p* < .001, *d* = 0.71 (*M*_FMHO_ = .56, *SD*_FMHO_ = .09) and in the FMLO condition, *t*(35) = 4.76, *p* < .001, *d* = 0.79 (*M*_FMLO_ = .58, *SD*_FMLO_ = .09). Exploratory post-hoc analyses showed that recognition accuracy did not differ between encoding conditions, *t*(35) = 0.54, *p* = .590, two-tailed.

### 2.3 Discussion

If a newly learned label is well integrated in neocortical networks, it is expected to compete with its lexical neighbors. Consequently, more time is required until these neighbors can be uniquely identified. In order to test lexical competition, we assessed response latencies to hermit words, expecting slowed responses to hermits which had artificially been assigned a new neighbor at encoding (neighbor hermit trials), compared to hermits which had not obtained a new lexical neighbor (non-neighbor hermit trials). Since Coutanche and Thompson-Schill (2014) had already shown that rapid lexical integration by means of FM but not EE is possible, we wanted to extend this research question and investigate if feature overlap might modulate rapid lexico-semantic integration of the picture-label associations within the FM encoding condition, using lexical competition as measure of integration. We observed a lexical competition effect already shortly after the labels had been encoded by means of FM. Consistent with Coutanche and Thompson-Schill’s (2014) results, our findings show that the labels of the novel associations were integrated immediately after FM encoding.

In contrast to our expectations, the lexical competition effect for FMHO trials was not different from that for FMLO trials, and numerically even smaller. We would like to offer an explanation for the lack of the expected moderating effect of feature overlap on lexical competition after learning by means of FM. Although we argued that the integration of the picture-label associations should provoke lexical competition, integration of the complete associations (i.e., the specific combination of the picture together with the label) likely is a sufficient but not a necessary condition to observe a lexical competition effect for the label. Contrary to what we proposed prior to the experiment, we now think that it might be more precise to say the lexical competition effect reflects lexical integration of the labels on an item level but not necessarily semantic integration of the labels with their associated pictures. Thus, lexical competition can be observed even though the complete associations are not integrated. The reason why we predicted a larger effect for FMHO trials was that stronger PrC recruitment in this encoding condition should have fostered especially semantic integration of the complete picture-label associations, due to increased PrC involvement in the discrimination of pictures sharing many features. Given that integration of the specific picture-label combination is not necessary to induce lexical competition, the absence of a feature overlap effect on lexical competition does not allow for conclusions about semantic integration of the complete associations. Furthermore, if our assumption holds true that manipulation of feature overlap reflects differential PrC involvement, it might not have been advantageous to manipulate feature overlap within subjects with trials of different overlap conditions presented in random order. It is likely that this could have severely decreased the signal-to-noise ratio, such that residual PrC activity of the previous trials might have blurred PrC activity of the current trial.

The semantic priming task we used as a measure of semantic integration in Experiment 2 should bring more clarity to the role of feature overlap in the integration of the complete associations. We manipulated feature overlap between subjects and extended the design with an EE condition.

## 3. Experiment 2

In Experiment 2, we administered a semantic priming task on two consecutive days. Since the semantic priming results reported by Coutanche and Thompson-Schill (2014) are difficult to interpret due to potential confounds, we considered it necessary to obtain comparable data not only for the different overlap conditions within FM encoding (as in Experiment 1) but also for an EE condition. As we used items of different semantic categories at encoding, it was possible to counterbalance for categories of the priming targets. In order to provoke stimulus processing on a more elaborate semantic level, we used a task requiring a semantic instead of a lexical decision. We predicted rapid semantic integration (i.e., a priming effect shortly after encoding) in the FMHO condition, and this effect should be larger than in the FMLO condition. We expected no priming effect in the EE condition on the same day but an increased priming effect after 24 hours as there should have been enough time for gradual consolidation into neocortical structures. For the FMLO condition, we did not predict a priming effect immediately after encoding as no use could be made of the catalyzing effect of feature overlap (as in the FMHO condition). It cannot be excluded that there might also be hippocampal engagement at encoding in the FM conditions in young and healthy participants, which potentially could foster semantic priming after 24 hours of consolidation. However, hippocampal involvement at FM encoding is presumably much less than in the EE condition. As no direct integration should have taken place in the FMLO condition and hippocampal contribution to learning should be negligible, we did not expect a semantic priming effect after FMLO encoding on Day 2.

In addition to these implicit measures of integration, we conducted a three-alternative forced-choice recognition test in order to examine if participants also showed explicit learning above chance level and if the EE group showed better recognition performance than the FM groups. This would be expected as healthy participants should benefit from intentional learning in the EE condition if tested with explicit recognition tests. Again, we did not make predictions on differences in recognition accuracy between the two FM conditions (FMHO, FMLO) since it cannot be disentangled to what extent retrieval is based on hippocampal or cortical processing.

### 3.1 Methods

#### 3.1.1 Participants

As encoding condition was manipulated between subjects, 120 participants were randomly allocated to one of three encoding conditions (FMHO, FMLO, EE). Four participants had to be excluded from all analyses as they had already taken part in another experiment in which the same stimulus material was used, leading to an overall sample size of *N* = 116 participants (*n*_FMHO_ = 39, *n*_FMLO_ = 39, *n*_EE_ = 38; 96 female; *M*_age_ = 23.1 years, age range: 18-35 years). There was no age difference between groups, *F* < 1. All participants were students from Saarland University, native German speakers, and had normal or corrected-to-normal vision. The experiment was split into two sessions of approximately 20-25 minutes each, separated by 24 hours (range: 23.4-24.4 hours). Participants gave written informed consent prior to the experiment on Day 1 and were compensated for their participation with 8€ per hour after completion of the experiment on Day 2. The experiment was approved by the local ethics committee of Saarland University in accordance with the declaration of Helsinki.

#### 3.1.2 Materials

Forty-eight pairs of picture pairs were drawn from the stimulus material of the previously conducted rating study (see Experiment 1). Only those item pairs were included for which the previously unknown item had been classified as unknown by most participants in the rating study (on average, by 88 %, *SD* = 12 %) and had been rated with the lowest familiarity (*M* = 2.01, *SD* = 0.42; 1 = *not at all familiar*, 5 = *very familiar*), and for which the previously known item had been rated as known by most participants (on average, by 90 %, *SD* = 13 %) and with the highest familiarity (*M* = 4.44, *SD* = 0.41). Moreover, only pairs of item pairs with the highest difference between the overlap rating of the high-overlap and the low-overlap item pair were included (*M*_FMHO_ = 3.62, *SD*_FMHO_ = 0.53; *M*_FMLO_ = 1.42, *SD*_FMLO_ = 0.39; 1 = *not at all similar*, 5 = *very similar*; *M*_diff_ = 2.20, *SD*_diff_ = 0.68). In the final item set, familiarity for the previously unknown items was significantly lower than for the previously known items, significantly more participants of the rating study had rated the previously known item as known than the previously unknown items, and overlap of the high feature overlap pairs was higher than overlap of the low feature overlap pairs (all *p*s < .001). In addition, the high overlap item pair with the lowest overlap still had a higher overlap rating than the low overlap item pair with the highest overlap. Further 16 trials were added as filler trials, of which two trials were presented as buffer trials each at the beginning and at the end of the encoding phase each. Filler trials matched the participants’ encoding condition and were excluded from all analyses. The size of the pictures varied depending on the items’ relative size in reality, but was 300 x 300 pixels at maximum, leading to a maximum visual angle of approximately 8.2°.

Labels remained the same as in Experiment 1. Those items which had been assigned three hermit neighbor labels for usage in the lexical competition task in Experiment 1 were given one of these three names. The labels used for Experiment 2 consisted of 4-9 letters with a mean length of *M* = 6.13 letters (*SD* = 1.18). In the two semantic priming phases, the labels of the previously unknown items were presented as primes, followed by a familiar German noun as target. Target words were either animals or plants. Each prime was assigned to four targets: two semantically related targets (same category as the prime) and two unrelated targets (different category). Unrelated prime-target pairs were created by reallocating targets to unrelated primes. All primes were presented twice, once on each day, whereas targets were only presented once. Within each participant, 25% of the primes were presented together with a related target only on Day 1, 25% only on Day 2, 25% on both days, and 25% on neither day. Assignment of trials to relatedness condition was counterbalanced across participants. Targets were of low frequency (lemma frequencies between 0.01 and 12.57 per million words; *M* = 1.82, *SD* = 2.48; Heister et al., 2011) and preferably long (4-13 letters; *M* = 7.33, *SD* = 1.89) as it has been shown that priming effects can be strengthened if processing of the target word takes participants more time (Hines, Czerwinski, Sawyer, & Dwyer, 1986). None of the targets had been presented previously in the experiment, neither as words nor as pictures of previously known items in the encoding phase. All prime and target words were displayed in the center of the screen in Arial 27 point font.

For the three-alternative forced-choice recognition test the target picture was paired with two foil pictures from the same higher-level category (either all plants or all animals). All pictures appeared three times (once as target, twice as foil), separated by at least four trials. Test foils had all been used as previously unknown items in the encoding phase in order to control for item familiarity. All other constraints were as in Experiment 1.

#### 3.1.3 Design and Procedure

##### Encoding

The experimental settings for all three groups (FMHO, FMLO, EE), the cover story, and the encoding procedure for the two FM groups were equal to Experiment 1. In contrast to the FM groups, learning was intentional in the EE group as this group was informed about the later associative recognition memory test. In the EE encoding phase, participants were only presented with the picture of the previously unknown item (see Figure 4). Contrary to previous studies (cf. Atir-Sharon et al., 2015; Coutanche & Thompson-Schill, 2014; Greve et al., 2014; Himmer et al., 2017; Korenic et al., 2016; Merhav et al., 2014, 2015; Sharon et al., 2011; Smith et al., 2014; Warren & Duff, 2014; Warren et al., 2016), the EE group was presented with the same questions as the FM groups, in order to prevent any confounds due to inconsistencies in task demands apart from the critical FM determinants. Before the actual experiment started, all three groups conducted a practice phase of six encoding trials. In addition to the 48 experimental encoding trials, 16 filler trials were inserted. In the filler trials, the question referred to the previously known item, of which two trials were inserted as buffer trials each at the beginning and at the end of the encoding phase. Stimulus presentation was as in Experiment 1.

**Figure 4.**
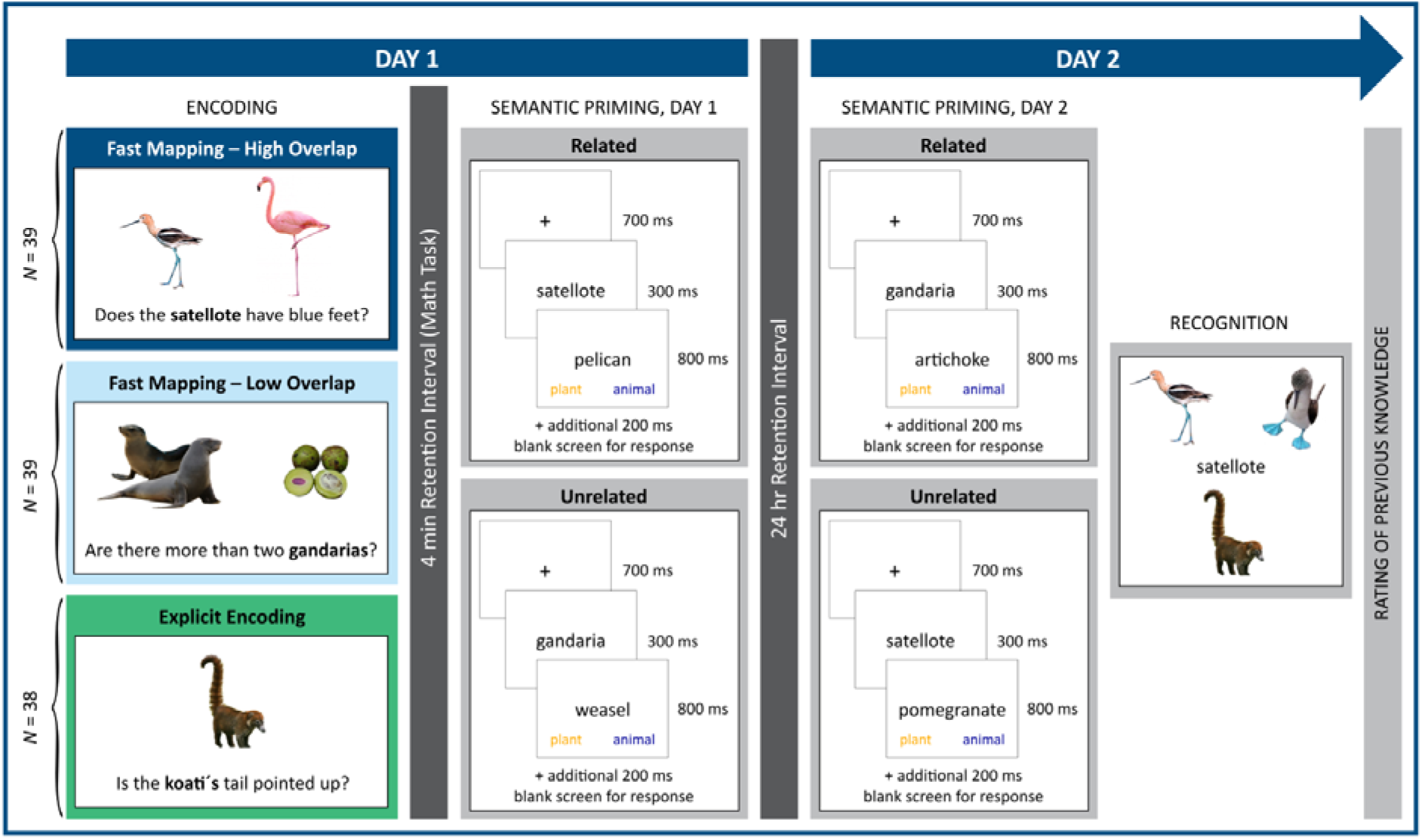
Experimental design and procedure of Experiment 2. **Encoding.** Encoding condition was manipulated between subjects. Participants in the explicit encoding condition were explicitly instructed to remember the item. After the question had been presented for 2000 ms, pictures were inserted and presented together with the question for 3500 ms. Response options (*yes*/*no*) were provided after both pictures and question had disappeared. Feedback was given after a response had been made. **Semantic priming.** For the semantic priming phases on Day 1 and Day 2, relatedness was fully counterbalanced across participants and study-test delays. **Recognition.** In the three-alternative forced-choice recognition test, the screen was presented as depicted in the figure for 3000 ms and then a prompt to respond appeared at the bottom of the screen. Targets and foils within one display always belonged to the same higher-level category (i.e., either both items were animals or both items were plants).

##### Semantic Priming

All following phases were identical for the three groups. After a 4-minute retention interval, in which participants solved simple mathematical equations, the first of two priming phases started. Both priming phases were preceded by a practice phase of six trials, in which primes were pseudo-words that had not appeared in the encoding phase. In order to accustom participants to the task demands, two buffer trials of the same nature as the practice trials were inserted at the beginning of each priming phase. Each trial began with the presentation of a fixation cross in the center of the screen for 700 ms, followed by a prime for 300 ms, which was the label of a previously unknown item of the encoding phase (see Figure 4). Next, the prime was replaced by the target, which was either semantically related or unrelated to the prime. The participants’ task was to indicate by keypress if the target was an animal or a plant, and as in Experiment 1, instructions emphasized speed over accuracy. Participants were informed that due to the fast pace of the task, they might make mistakes but nevertheless should focus on responding as fast as possible (as recommended by Wentura & Degner, 2010). The words *animal* and *plant* were displayed in blue and orange color on the left and right side on the bottom of the screen (color and position counterbalanced between subjects). Targets remained on the screen until participants responded by pressing the respective orange or blue key on the computer keyboard but for 800 ms at maximum. If no key had been pressed within 800 ms of target presentation, a blank screen was inserted for additional 200 ms in which the target was not visible but responses were still recorded. All stimuli of the priming phase were presented in random order in the center of the screen and displayed in Arial 27 point font. After a delay of 24 hours, a second priming phase was administered, in which the same primes were presented as on Day 1 but together with different targets. Apart from that, the procedure was kept identical with the Day 1 priming phase.

##### Recognition

After the completion of the second priming phase on Day 2, a three-alternative forced-choice recognition test was administered. A fixation cross was displayed in the center of the screen for 500 ms, before it was replaced by the recognition test label in Arial 27 point font (see Figure 4). The test target picture and the two test foil pictures were arranged around the label, with their positions on the screen randomly assigned (top-left, top-right, bottom-center). Participants were instructed to indicate which of the three pictures belonged to the test label by clicking on the respective picture. In order to ensure that all participants had enough time to thoroughly look at all three pictures, responses could not be given before 3000 ms of stimulus presentation, after which a verbal prompt to respond appeared at the bottom of the screen. As soon as a decision was made, the next trial started and the mouse cursor was automatically set back to the center of the screen. If no key had been pressed within 6000 ms of stimulus presentation, participants were encouraged to respond faster and the next trial started.

##### Rating of previous knowledge

At last, participants rated how well they had known the items prior to the experiment. Rating instructions and procedure were identical to Experiment 1, except that a 6-point Likert scale was used (1 = *had not known the item at all before the experiment*; 6 = *had known the item very well before the experiment*). If a rating of >= 4 was given, participants were asked to type in the item’s name at the lowest category level possible.

#### 3.1.4 Data analysis

The semantic priming effect was calculated by subtracting response times for correct responses to related targets from response times for correct responses to unrelated targets, individually for each participant. Analyses included all correct trials for which the individual ratings of both the known and the unknown item (EE: only the unknown item) were congruent with the expected knowledge, that is, items classified as unknown in the rating study with an individual knowledge rating of <= 3, and items previously classified as known with a rating of >= 4 (mean dropout rate was 7.7% for both days). Further trials were excluded if response latencies were 1.5 interquartile ranges below the first quartile or above the third quartile of individual response times (Tukey, 1977).

For Day 1 analyses, nine participants had to be excluded because they had not performed above chance level in the semantic priming task (2 participants of the FMHO group, 3 FMLO, 1 EE; *p* > .05, binomial test) or were outliers with regard to the semantic priming effect according to Tukey (1977; 1 FMLO, 2 EE), resulting in an overall sample size of *N*_Day1_ = 107 (*n*_FMHO_ = 37, *n*_FMLO_ = 35, *n*_EE_ = 35). Participants who were classified as outliers with regard to the priming effect were again included in Day 2 analyses whereas chance performers were excluded from all further analyses as we took low performance in such an easy task as an indicator of a lack of motivation and subsequent performance would likely be based on less overall attendance to the stimuli.

In addition to the chance performers of Day 1, two more participants were excluded for the same reason on Day 2 (1 FMLO, 1 EE). Four participants were outliers regarding the priming effect on Day 2 (4 FMHO), resulting in an overall sample size of *N*_Day2_ = 104 (*n*_FMHO_ = 33, *n*_FMLO_ = 35, *n*_EE_ = 36). For the recognition test, only participants who performed at chance in at least one priming phase were excluded (*N*_rec_ = 108; *n*_FMHO_ = 37, *n*_FMLO_ = 35, *n*_EE_ = 36). Recognition accuracy represents proportion of correct responses.

If not noted differently, *t* tests to compare semantic priming effects were one-tailed and the significance level was set to α = .05. Effect size *d* for the between-subjects comparisons of the semantic priming effect was calculated as difference of the mean semantic priming effects, divided by the pooled standard deviation of the effects. All other analyses remained the same as in Experiment 1.

### 3.2 Results

#### Semantic Priming, Day 1

Accuracies in the semantic priming task were above chance in all encoding conditions (all *p*s < .001; see Table 2 for accuracies). A one-way ANOVA with the between-subjects factor encoding condition (FMHO, FMLO, EE) did not show a significant effect of encoding condition on the semantic priming effect, *F*(2,104) = 2.14, *p* = .123. As we were especially interested in the comparison of the FM groups, we additionally investigated the differences between FMHO and FMLO. In line with our hypotheses, planned comparisons showed that the semantic priming effect was significantly larger for the FMHO group than for the FMLO group, *t*(70) = 1.96, *p* = .027, *d* = 0.46, although the difference between the FMHO and the EE group was not significant, *t*(70) = 1.09, *p* = .140 (see Figure 5; see Table 2 for response times). There was a significant semantic priming effect in the FMHO condition, *t*(36) = 1.72, *p* = .047, *d* = 0.28 (*M* = 9.57 ms, *SD* = 33.79 ms), but neither in the FMLO condition, *t*(34) = −1.07, *p* = .294, two-tailed (*M* = −6.33 ms, *SD* = 35.17), nor in the EE condition, *t*(34) = 0.32, *p* = .749, two-tailed (*M* = 1.55 ms, *SD* = 28.40). If the semantic priming effect after FM encoding was calculated across overlap conditions, no priming effect was found, *t* < 1.

**Table 2.**
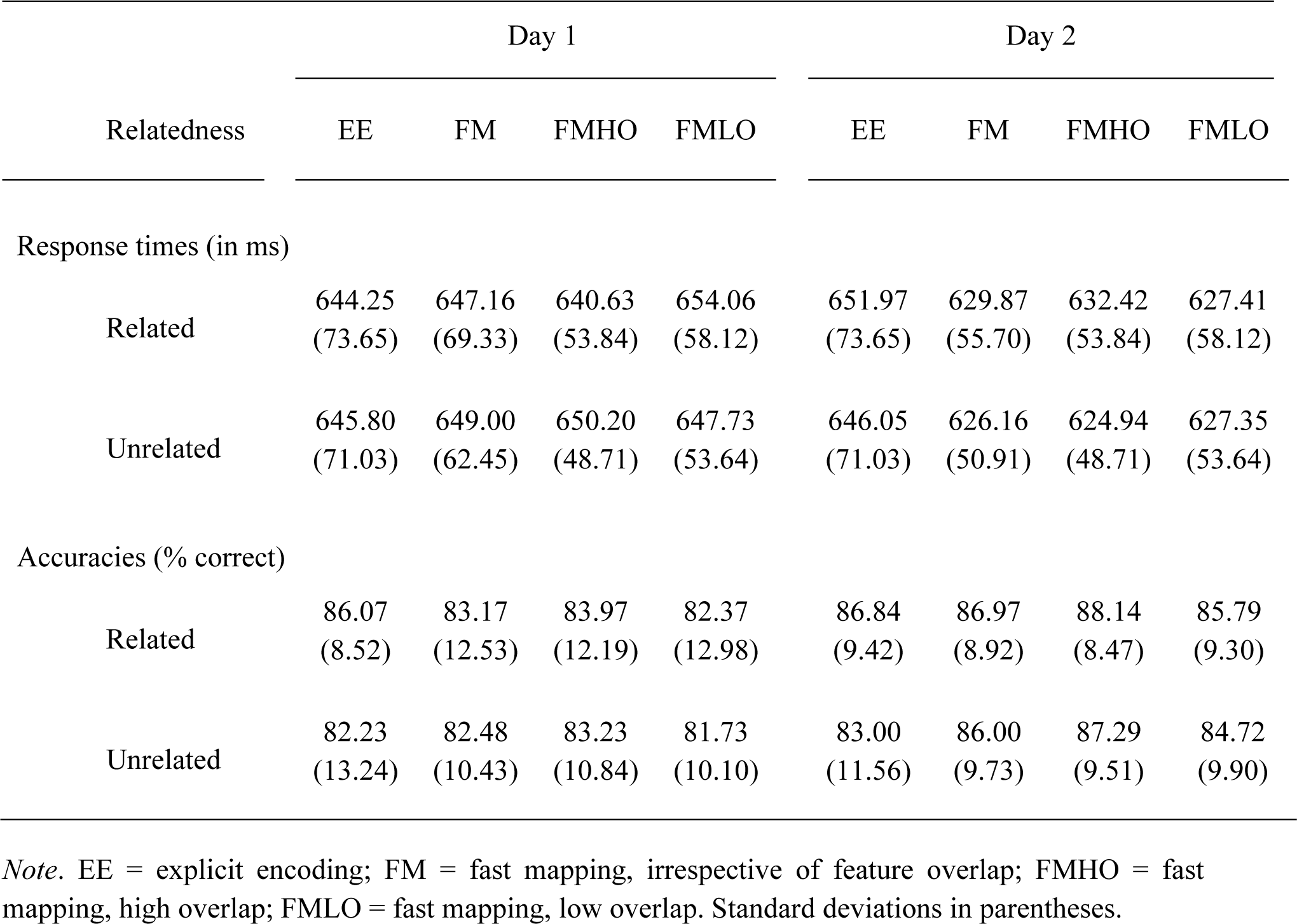
Mean Response Times (in ms) and Mean Accuracies (in % Correct) by Relatedness and Encoding Condition in the Semantic Priming Task of Experiment 2, separately for Day 1 and Day 2

**Figure 5.**
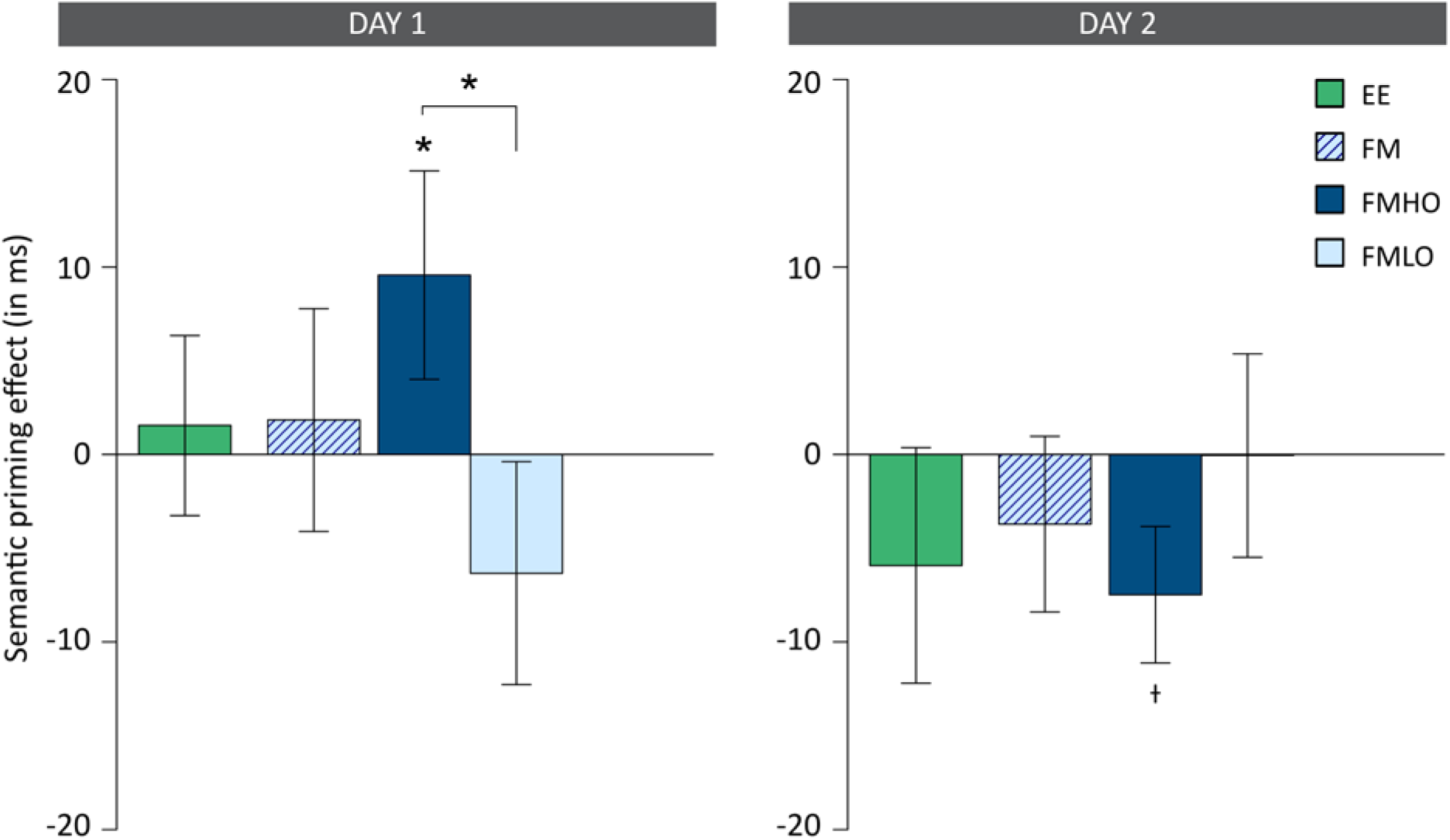
Results of the semantic priming task of Experiment 2 for Day 1 and Day 2. The semantic priming effect was calculated by subtracting response times to related targets from response times to unrelated targets. Error bars represent the standard error of the means. EE = explicit encoding; FM = fast mapping, irrespective of feature overlap; FMHO = fast mapping, high overlap; FMLO = fast mapping, low overlap. †*p* < .10, \**p* < .05

#### Semantic Priming, Day 2

Accuracies in the semantic priming task were above chance in all encoding conditions (all *p*s < .001; see Table 2 for accuracies). For the Day 2 semantic priming task, the one-way between-subjects ANOVA again did not reveal a significant effect of encoding condition on the semantic priming effect, *F* < 1 (Figure 5; see Table 2 for response times). Contrary to our hypotheses, post-hoc *t* tests revealed a numeric tendency towards a negative semantic priming effect for the FMHO group and the EE group. As there is literature on the phenomenon of negative priming effects which might be interesting in the context of learning by means of FM, we wanted to further investigate if this negative tendency deviated significantly from zero, clearly stressing that these analyses were calculated post hoc and are to be considered as exploratory only. These two-tailed post-hoc *t*-tests revealed a marginally significant negative semantic priming effect, *t*(33) = −1.97, *p* = .057, two-tailed, *d* = 0.39 (*M* = −7.48 ms, *SD* = 22.15), for the FMHO group and again no significant priming effect for neither the FMLO group (*M* = −0.05 ms, *SD* = 32.10) nor the EE group (*M* = −5.92 ms, *SD* = 37.16), both *t*s < 1, two-tailed. If the semantic priming effect after FM encoding was calculated across overlap conditions, no priming effect was found, *t*(68) = −1.11, *p* = .270.

#### Recognition

In the three-alternative forced-choice recognition test, which was conducted after the semantic priming phase on Day 2, all groups performed above chance level (all *p*s < .001). A one-way between-subjects ANOVA of encoding condition on recognition accuracy did not reach significance, *F*(2,105) = 2.23, *p* = .113. As expected, accuracy of the EE group was superior to accuracy of the FM groups, *t*(106) = 1.67, *p* = .049, *d* = 0.40 (*M*_EE_ = .52, *SD*_EE_ = .13; *M*_FM_ = .48, *SD*_FM_ = .08; *M*_FMHO_ = .50, *SD*_FMHO_ = .08; *M*_FMLO_ = .47, *SD*_FMLO_ = .08).

### 3.3 Discussion

When the semantic priming effects of Experiment 2 are calculated across both FM groups, we did not observe semantic priming for the FM condition, neither on Day 1 nor on Day 2. Strikingly, when the FMHO and FMLO groups were analyzed separately, we found a semantic priming effect for the FMHO group on Day 1, but neither for the FMLO group nor for the EE group. Moreover, the priming effects in the FMHO group and in the FMLO group were significantly different. This indicates that rapid semantic integration of novel associations by means of FM is possible, but only if feature overlap between the previously known and the previously unknown item is high, leading to better integration of the picture-label association. Although the semantic priming effect for the FMHO group did not significantly differ from the priming effect for the EE group, a semantic priming effect for associations encoded through EE immediately after encoding was not observed, indicating that there was no rapid semantic integration after encoding by means of EE.

Whereas the lack of a priming effect in the FMLO group on Day 2 hat been predicted, the expected semantic priming effect for the EE group on Day 2 was not found. It is conceivable that consolidation processes might possibly have been overshadowed by a weakening of the associations overnight. Despite better integration of the associations after 24 hours, explicit retrieval might still have become too effortful after a longer delay, especially considering that, contrary to other studies (e.g., Coutanche & Thompson-Schill, 2014; Greve et al., 2014; Merhav et al., 2014, 2015; Sharon et al., 2011; Warren & Duff, 2014; Warren et al., 2016), participants encoded the associations only once. This is additionally evident in rather weak recognition accuracy levels (which were also assessed on Day 2), compared to recognition performance typically found in EE learning, which might also be due to more effortful encoding task requirements. Whereas participants are typically only instructed to remember the depicted item in the EE condition, we additionally asked them to answer the same question as in the FM condition.

Contrary to our expectations, exploratory post-hoc analyses revealed a marginally significant negative semantic priming effect for the FMHO group on Day 2. Although this marginally significant negativity should only be interpreted very carefully as it was not expected, we would like to provide an explanation for such a tendency as negative priming effects are a not an uncommon phenomenon and we do not want to leave this unexpected pattern uncommented. In previous literature on semantic priming, negative priming effects are explained by a center-surround approach (Walley & Weiden, 1973; see also Carr & Dagenbach, 1990). According to this approach, strongly related targets are inhibited if access to the primes is weak, thereby leading to prolonged response latencies for related targets. In line with this, it is conceivable that primes in the FMHO condition were more difficult to access after 24 hours. We want to clearly point out that this is only a post-hoc explanation trying to address the numerically strong negativity but should not be over-interpreted.

## 4. General Discussion

It has been proposed that FM might be a learning paradigm that allows for rapid, direct integration of novel associations (e.g., Atir-Sharon et al., 2015; Coutanche & Thompson-Schill, 2014; Himmer et al., 2017; Merhav et al., 2014, 2015; Sharon et al., 2011). Yet, contradictory findings have been reported (e.g., Greve et al., 2014; Smith et al., 2014; Warren & Duff, 2014; Warren et al., 2016) and it remains to be clarified which factors could possibly moderate the learning benefit of FM.

In Experiment 1, we observed a general lexical competition effect across feature overlap conditions, indicating that lexical integration by means of FM is generally possible. This is also consistent with the findings by Coutanche and Thompson-Schill (2014), who additionally showed that lexical integration was only found for an FM condition but neither for an EE condition nor for incidental learning per se. Here, we set out to further examine the essential criteria for learning success within the FM condition. Unexpectedly, the lexical competition effect we found was not larger in the FMHO condition than in the FMLO condition. This might have been due to the nature of the task as the paradigm we used only captures lexical integration of the label, which might be independent of semantic integration of the complete picture-label combinations. Thus, we might only have captured lexical integration of the labels on an item level but not lexico-semantic integration of the complete associations. We think that it is conceivable that the FM paradigm as defined by Sharon et al. (2011) is not only appropriate to evoke immediate semantic integration of the associations but also lexical integration of the labels only, irrespective of feature overlap, although it has been shown that at least the presence of two pictures is a necessary determinant for rapid lexical integration (Coutanche & Thompson-Schill, 2014). In a more recent study, Coutanche and Koch (2017) further investigated the role of the previously known item at encoding. Using a similar design as Coutanche and Thompson-Schill (2014), they observed lexical competition after FM encoding but only if the previously known items were atypical for their category. In our experiments, the previously known items were counterbalanced across conditions and thus typicality was kept constant. Hence, the lack of an effect of feature overlap on lexical competition in our Experiment 1 does not contradict their effect of typicality. Whereas typicality of the previously known items might affect lexical integration, visual discrimination and in particular the degree of feature overlap does not, as shown in our Experiment 1. Further research is needed in order to identify potentially differential effects of feature overlap and typicality of the previously known item.

In Experiment 2, we found a semantic priming effect immediately after learning for the FMHO group but neither for the FMLO group nor the EE group, indicating that rapid semantic integration of the complete picture-label associations is possible in an FM paradigm, but only if the unknown and the known item share many features. This might be due to higher PrC involvement in the FMHO condition, although these behavioral data do not allow for interpretations on a neurofunctional level.

### 4.1 Potential underlying neurocognitive mechanisms

Although we do not draw conclusions from our behavioral findings on underlying neurocognitive mechanisms, we originally approached the identification of factors potentially moderating rapid learning by means of FM from a neurocognitive perspective. The functional and representational characteristics of the PrC, that is, its involvement in recognition memory and perceptual and conceptual processing of higher-order object representations, let assume that it is especially qualified to support learning by means or FM. We manipulated feature overlap between the previously known and the previously unknown item in the FM encoding phase, based on the idea that PrC recruitment is associated with the discrimination of items sharing many features. Whereas most previous studies point to the ATL as candidate structure to be mainly involved in learning by means of FM, we do not assume that the implementation of neocortical integration in the FM paradigm is restricted to either the ATL or the PrC but rather suggest that both structures might be highly involved. Ranganath and Ritchey (2012) even consider the PrC a core component of an anterior temporal system that also includes the ATL. Whereas the ATL is ascribed the integration of features of various modalities from their respective cortical sensory areas to a coherent whole (see Lambon Ralph et al., 2017, for a review), the PrC is considered especially responsible for the distinct identification of unique cross-modal feature combinations and their discrimination from similar objects (e.g., Kivisaari, Monsch, & Taylor, 2013; Kivisaari, Tyler, Monsch, & Taylor, 2012; Taylor, Moss, Stamatakis, & Tyler, 2006; Tyler et al., 2004). With regard to learning by means of FM, it is conceivable that ATL engagement might increase when the previously known item is atypical for its category (see Coutanche & Koch, 2017) and that PrC engagement might increase when the known and the unknown item share many features. However, this remains to be further investigated in brain imaging studies.

### 4.2 Visual or semantic overlap?

We primarily defined feature overlap as visual similarity between two items, that is, the number of (visual) features they have in common. However, in doing so, we might inevitably have manipulated semantic similarity as well, as highly similar looking objects usually also belong to the same semantic category. There would be no need to disentangle visual and semantic overlap if both addressed the same underlying process. However, it is conceivable that whereas the manipulation of *visual* overlap in the FM paradigm most likely affected the initial visual *discrimination* between pictures, the simultaneous manipulation of *semantic* overlap might, in a next step, have influenced semantic *integration*. In order to disentangle the effects of visual and semantic overlap in our data pattern, we conducted further post-hoc analyses, although it needs to be said that these do not allow for final conclusions on the exact contributions of each process. The discrimination between highly overlapping items in our experiments always required a decision within the same lower-level category (e.g., both were birds), whereas low-overlap item pairs were always from different lower-level categories but sometimes from the same higher-level category (e.g., both animals: a bird and a mammal) and sometimes from different higher-level categories (e.g., an animal and a plant; see Figure 1). In order to further examine if semantic overlap is the crucial factor on which the difference between the FMHO and FMLO priming effects is based, we calculated the semantic priming effect post-hoc for the FMLO condition of Day 1 separately for FMLO pairs from the same higher-level category and for FMLO pairs from different higher-level categories. These analyses revealed that the semantic priming effect was not different between FMLO pairs of the same higher-level category and those of different higher-level categories, *t* < 1. Moreover, if the difference between the semantic priming effects of the FMHO and FMLO group was simply based on semantic overlap, this difference should disappear if only FMLO item pairs of the same higher-level category were considered for the analyses. Although the comparison between the semantic priming effect of the FMHO group and the FMLO group showed only a trend towards significance after the exclusion of FMLO trials using pairs of different higher-level categories, *t*(71) = 1.28, *p* = .102, one-tailed, the pattern remained the same, as the priming effect for FMLO trials of the same higher-level category (*M* = −4.04 ms, *SD* = 52.05 ms) still was on a similar level as the priming effect for all trials of the FMLO group (*M* = −6.33 ms, *SD* = 35.17). We therefore suggest that visual feature overlap between the unknown and the known item decisively drives the difference between the priming effect in the FMHO and the FMLO condition although we would assume that both visual and semantic overlap are essential for successful learning by means of FM. However, valid conclusions on the contribution of visual versus semantic overlap would require an a-priori manipulation of both processes separately.

### 4.3 Stability of memory representations acquired by means of FM

Previous notions in the literature often emphasize that memories acquired by means of FM are maintained over time, based on the finding that recognition test performance remains above chance even after longer delays (e.g., Coutanche & Thompson-Schill, 2014; Greve et al., 2014; Korenic et al., 2016; Merhav et al., 2015; Sharon et al., 2011; but see Smith et al., 2014). However, it is difficult to draw general conclusions on the robustness of memory representations in FM learning from the present literature. There is a great variety of study-test delays, regarding the duration of the delay (from no delay to a one-week delay), the nature of the filler task and its level of interference (e.g., a vocabulary test, Sharon et al., 2011, and Coutanche & Thompson-Schill, 2014; conversation, Smith et al., 2014; an intelligence test, Greve et al., 2014; math tasks in our experiments), and potential carry-over effects through other (memory) tests that were conducted between the encoding and recognition phase (e.g., free recall of the associations prior to the recognition test, Warren & Duff, 2014, and Warren et al., 2016). In addition, accuracy in explicit forced-choice recognition tests might not be an appropriate measure to investigate robustness in a longitudinal design. Repeated explicit testing within participants inevitably adds noise to measures of neocortical integration and hence, test accuracy no longer represents pure incidental FM learning. We consider repetition of implicit measures, such as the semantic priming task in Experiment 2, to be less critical, since the newly learned associations have never been explicitly retrieved before the recognition test, which was administered only once and after all semantic priming tasks had been completed. At a first glance, our finding that the semantic priming effect on Day 1 for the FMHO group in Experiment 2 disappeared and even (numerically) turned into a negative direction on Day 2 seems to contradict the assumption of stability of memories acquired by means of FM. However, a potential explanation could be that access to the newly acquired labels (which were presented as primes in the semantic priming task) might have been weakened overnight, leading to lateral inhibition of semantically related items in order to suppress interference while accessing the weaker memories. Therefore, a stable association between prime and target after one day could be reflected in prolonged response times to the targets if the primes are only weakly accessible (see e.g., Carr & Dagenbach, 1990).

### 4.4 Limitations

In our experiments, the number of trials was higher than in most FM experiments (92 trials in Experiment 1 and 48 trials in Experiment 2, instead of 16-24 trials per encoding condition, see e.g., Coutanche & Thompson-Schill, 2014; Greve et al., 2014; Sharon et al., 2011; but see also Merhav et al., 2015, who used 50 trials per encoding condition). This could have led to more interference already during encoding, thereby impeding the integration of the associations. Moreover, using more trials makes it difficult to provide a stimulus list consisting of heterogeneous materials. Homogeneity of the stimulus material could have increased this interference, especially for unfamiliar items (see Brandt, Zaiser, & Schnuerch, 2018). Apart from that, questions were only presented visually instead of bimodally as in other FM studies (e.g., Greve et al., 2014; Merhav et al., 2014; Sharon et al., 2011; Smith et al., 2014) and, in contrast to other studies (see above), each association was encoded only once. We nevertheless think such a single-exposure encoding procedure is suited best to investigate differential effects between encoding conditions as in that way, they can clearly be attributed to differences in pure encoding processes and cannot be influenced by retrieval processes during repeated presentations at encoding. These deviations of the encoding phase could have led to smaller effects of integration than we might have observed if associations were encoded on a deeper level. It is noticeable, however, that the results of both experiments clearly revealed immediate integration of novel associations through FM encoding, despite relatively high numbers of trials and, most importantly, even after only a single exposure to the associations.

## 5. Conclusions

Our findings provide further evidence for rapid integration of novel, arbitrary picture-label associations by means of FM. Integration of the newly learned labels led to both lexical competition and semantic priming immediately after encoding. Whereas lexical competition was unaffected by feature overlap in the FM encoding phase, an immediate semantic priming effect was only found if the items shared many features, implying that a high feature overlap is essential for semantic integration of novel picture-label associations. Evidence for rapid semantic integration has not yet been observed and with our findings we can provide explanations of results previously reported in the literature. However, the underlying mechanisms of these findings yet need to be identified. As we cannot draw conclusions on the role of PrC involvement in learning by means of FM with our behavioral results, this remains to be further investigated on a neurofunctional level.

## Acknowledgments

This work was funded by the German Research Foundation [grant numbers BA 5381/1-1 and ME 4484/1-1] in partial fulfillment of A.-K. Z’s doctoral dissertation. We would like to thank Alexander Hauck for his help with stimulus preparation and data collection.

## Appendix

Newly Created Lexical Neighbors to German Hermit Words, Used as Labels in Experiment 1 in Order to Evoke Lexical Competition

| Neighbor | Hermit | Translation | Neighbor | Hermit | Translation |
| --- | --- | --- | --- | --- | --- |
| *Akroyat* | Akrobat | acrobat | *Matralle* | Matratze | mattress |
| *Albur* | Album | album | *Menka* | Mensa | canteen |
| *Borbe* | Bombe | bomb | *Minuster* | Minister | minister |
| *Brude* | Bruder | brother | *Mored* | Moped | moped |
| *Dontor* | Doktor | doctor | *Murtel* | Murmel | marble |
| *Eigel* | Eigelb | egg yolk | *Muspel* | Muskel | muscle |
| *Fabrek* | Fabrik | factory | *Orfel* | Orgel | pipe organ |
| *Famolie* | Familie | family | *Palist* | Palast | palace |
| *Flemme* | Flamme | flame | *Pfalle* | Pfanne | pan |
| *Förser* | Förster | forester | *Pilor* | Pilot | pilot |
| *Futo* | Foto | photo | *Pistoke* | Pistole | pistol |
| *Galanie* | Galaxie | galaxy | *Plakal* | Plakat | placard |
| *Globuk* | Globus | globus | *Satellot* | Satellit | satellite |
| *Gürmel* | Gürtel | belt | *Schirk* | Schirm | umbrella |
| *Honil* | Honig | honey | *Schneel* | Schnee | snow |
| *Kaisek* | Kaiser | emperor | *Stiemel* | Stiefel | boot |
| *Kalunder* | Kalender | calendar | *Taifur* | Taifun | typhoon |
| *Keramuk* | Keramik | ceramic | *Torado* | Tornado | tornado |
| *Kleiser* | Kleister | paste | *Trator* | Traktor | tractor |
| *Knoske* | Knospe | bud | *Trelor* | Tresor | safe |
| *Kondimor* | Konditor | confectioner | *Trochel* | Trommel | drum |
| *Künsler* | Künstler | artist | *Tunnek* | Tunnel | tunnel |
| *Lössel* | Löffel | spoon | *Tursine* | Turbine | turbine |
| *Magalin* | Magazin | magazine | *Vulka* | Vulkan | volcano |

*Note.* Participants were explicitly instructed to categorize persons or professions as *natural*. In the encoding phase, one third of the hermit neighbors were presented as labels in the FMHO (fast mapping, high overlap) condition, one third in the FMLO (fast mapping, low overlap) condition, and one third was not encoded, as in the lexical competition task, the respective hermit words served as non-neighbor hermits (i.e., hermit words which did not obtain a new neighbor at encoding).

## Footnotes

1 This does not necessarily mean that learning by means of FM is always hippocampus-independent. It has been shown that the hippocampus contributes to FM learning in healthy young adults (Atir-Sharon, Gilboa, Hazan, Koilis, & Manevitz, 2015) or at least it cannot finally be excluded that it is not involved (Merhav et al., 2015). We propose that in patients with severe and selective hippocampal lesions it is valid to conclude that hippocampal activity cannot have contributed to FM learning or retrieval. This does not account if functional integrity of the hippocampus is given.

